# Developing a rapid and highly efficient cowpea regeneration and transformation system using embryonic axis explants

**DOI:** 10.1101/738971

**Authors:** Ping Che, Shujun Chang, Marissa K. Simon, Zhifen Zhang, Ahmed Shaharyar, Jesse Ourada, Dennis O’Neill, Mijael Torres-Mendoza, Yinping Guo, Kathleen M. Marasigan, Jean-Philippe Vielle-Calzada, Peggy Ozias-Akins, Marc C. Albertsen, Todd J. Jones

## Abstract

Cowpea is one of the most important legume crops planted worldwide, especially in Sub-Saharan Africa and Asia. Despite decades of effort, genetic engineering of cowpea is still challenging due to inefficient *in vitro* shoot regeneration, *Agrobacterium*-mediated T-DNA delivery and transgenic selection. Here, we report a rapid and highly efficient cowpea transformation system using embryonic axis explants isolated from imbibed mature seeds. We found that removal of the shoot apical meristem by cutting through the middle of the epicotyl stimulated direct multiple shoot organogenesis from the cotyledonary node tissue. Furthermore, the application of a ternary transformation vector system using an optimized pVIR accessory plasmid provided high levels of *Agrobacterium*-mediated gene delivery. The utilization of spectinomycin as the selection agent enabled more efficient transgenic selection and plant recovery. Transgenic cowpea shoots developed exclusively from the cotyledonary nodes at high frequencies of 4.5 to 37% across a wide range of cowpea genotypes. We believe that the transformation principles established in this study could also be applied to other legumes to increase transformation efficiencies.

## Introduction

Domesticated in Africa and widely cultivated in the tropical and subtropical zones of the world, cowpea (*Vigna unguiculata* (L.) Walp.), also known as black-eyed pea, is one of the most valuable grain legumes for high-quality dietary protein, carbohydrates, lipids, minerals and vitamins for people in developing countries of Africa and Asia (Abdu Sani *et al*., 2015; Phillips *et al*., 2003; Singh, 2014). It is estimated that over 200 million people consume cowpea daily in Africa (Phillips *et al*., 2003; Singh, 2014). Despite its high tolerance to heat, dry conditions and soil acidity, cowpea is highly susceptible to insect pest and pathogen infestations resulting in lower productivity (Abdu Sani *et al*., 2015; Boukar *et al*., 2016; Obembe, 2008; Singh, 2014; Solleti *et al*., 2008a). Due to limited genetic variability of cowpea and strong cross-incompatibility between wild *Vigna* species and cultivated cowpea, little progress has been made in genetic improvement through conventional breeding to achieve desirable agronomic traits (Abdu Sani *et al*., 2015; Fang *et al*., 2007; Gomathinayagam *et al*., 1998; Latunde-Dada, 1990; Wamalwa *et al*., 2016). Hence, plant biotechnology provides an alternative approach to overcome those production constraints for improving the agronomic performance and developing better cowpea cultivars with higher grain quality and yield (Carlos Popelka *et al*., 2004; Zaidi *et al*., 2005). The development of insect-resistant cowpea, unsuccessful through conventional breeding, was successfully achieved by introducing *Bt* genes through genetic transformation and is a good example of plant biotechnology application in an orphan crop (Bakshi *et al*., 2011; Bett *et al*., 2017; Zaidi *et al*., 2005). Recently, significant progress has been made establishing genomic and gene expression data resources for two cowpea varieties, IT86D-1010 (Spriggs *et al*., 2018) and IT97K-499-35 (Lonardi *et al*., 2019; Munoz-Amatriain *et al*., 2017; Yao *et al*., 2016). However, the absence of an efficient genetic transformation system (Popelka *et al*., 2006; Somers *et al*., 2003) has impeded the full utilization of these resources for cowpea functional genomic studies to elucidate the mechanisms of heat and drought stress tolerance and to improve the agronomic traits, such as insect and pathogen resistances and increased productivity.

Legumes, especially cowpea, are known to be recalcitrant for genetic manipulation (Manman *et al*., 2013; Popelka *et al*., 2006; Solleti *et al*., 2008b; Somers *et al*., 2003). This is due to the absence of an amenable *in vitro* shoot regeneration system, the inadequate *Agrobacterium*-mediated T-DNA delivery to the targeted tissue and the inefficient transgenic selection methods for viable transgenic plant recovery. Therefore, published frequencies of *Agrobacterium*-mediated transformation are lower than 3.9% (Bett *et al*., 2019; Chaudhury *et al*., 2007; Manman *et al*., 2013; Mellor *et al*., 2012) and the process often requires more than 5 to 8 months (Chaudhury *et al*., 2007; Popelka *et al*., 2006). To overcome those obstacles and improve cowpea transformation efficiency, we evaluated shoot regeneration using embryonic axis (EA) explants isolated from imbibed mature seeds and identified that only cotyledonary node (cot-node) cells of the EA explants undergo rapid cell division and dedifferentiation to acquire organogenetic competence for shoot regeneration. Based on that observation, a rapid and highly efficient *in vitro* adventitious shoot regeneration system using EA explants was developed for direct *de novo* shoot organogenesis.

Next, we demonstrated that a ternary vector system developed previously (Anand *et al*., 2018) provided enhanced gene delivery not only for corn and sorghum transformation as we previously demonstrated (Anand *et al*., 2018; Che *et al*., 2018), but also for cowpea transformation using an auxotrophic *Agrobacterium* strain LBA4404 Thy-(thymidine mutant), a strain that reduces bacterial overgrowth during tissue culture compared to wild-type LBA4404. This ternary vector system contains the T-DNA binary vector and the optimized pVIR accessory (pPHP71539) plasmid with additional *vir* (Virulence) genes (Anand *et al*., 2018). The pVIR accessory plasmid and T-DNA binary vector have many desirable features, including smaller vector sizes, enhanced vector stability and amended *vir* genes for enhanced T-DNA gene delivery and ultimately higher transformation efficiency for all crops tested (Anand *et al*., 2018; Che *et al*., 2018).

To identify an optimal selection system for the EA-based *de novo* organogenesis, we also tested and compared the selection efficiency of *chloroplast transit peptide (CTP)-NPTII*/kanamycin (kan), *CTP-NPTII*/G418 and *CTP-spcN* (GenBank Accession No. AAD50455)/spectinomycin (spec) (Anada *et al*., 2017) selectable marker systems based on selection stringency and transgenic plant recovery. We found that the utilization of *CTP-spcN* combined with spec as the selectable marker facilitated the recovery of transgenic cowpea and provided more efficient and stringent transgenic selection and identification through the transformation process.

Finally, in order to test the robustness and flexibility of the protocol, an alternate spec selection system, *CTP-aadA*/spec (Martinell *et al*., 2017), and a hypervirulent *Agrobacterium* strain AGL1 (Lazo *et al*., 1991), which was previously used in cowpea transformation (Popelka *et al*., 2006), were utilized to test the transformability of nine cowpea genotypes for transgenic shoot regeneration and demonstrated the broad application potential of the transformation system.

Overall, we developed a rapid, robust and highly efficient *Agrobacterium*-mediated EA-based cowpea regeneration, transformation and selection system for generating transgenic cowpea exclusively from the cot-nodes of EA explants at a high frequency, between 4.5 to 37%, across a wide range of cowpea genotypes.

## Results and discussion

### De novo shoot organogenesis using EA as explants

A rapid, efficient and reproducible regeneration system is a prerequisite for establishment of an efficient cowpea genetic transformation system. Although several studies of *in vitro* regeneration of cowpea based on organogenesis have been reported (Aasim *et al*., 2010; Abdu Sani *et al*., 2015; Mamadou *et al*., 2008; Manman *et al*., 2013; Odutayo *et al*., 2005; Raveendar *et al*., 2009; Sani *et al*., 2018; Tie *et al*., 2013; Yusuf *et al*., 2008), an efficient cowpea regeneration system that enables highly efficient transformation is still lacking (Manman *et al*., 2013). Soybean transformation based on the preexisting meristems of EA explants has been well established and provides a reliable and highly efficient mean for introducing transgenes (Aragão *et al*., 2000; Liu *et al*., 2004; Turlapati *et al*., 2008). To test the regeneration efficiency of EA explants in cowpea, EA explants were isolated from imbibed mature seeds of cowpea variety IT86D-1010 by excising the cotyledons at the nodal points (Figure 1a). Those EA explants with the plumule excised (Figure 1b) were then cultured directly onto shoot induction medium (SIM) (Table S5) without selection in a vertical upright position with roots embedded in the media to induce shoot development. In most of the cases (> 95%), a single primary shoot was developed per EA when the shoot apical misterm (SAM) of the EA explants was kept intact (non-decapitated EA explants) during regeneration (Figure 1c). However, multiple shoot development was observed occasionally for a small number of explants (< 5%) (Figure 1d). Compared to the morphology of EA explants with single primary shoot development (Figure 1c), the EA explants with multiple shoots, regenerated exclusively around cot-nodes (Figure 1d), were much shorter and lacked both epicotyls and SAMs. The lack of epicotyls and SAMs could be due to the accidental damage of those tissues during EA isolation and plumule excision. This finding indicated that the *de novo* organogenesis of shoots around the cot-nodes could be inhibited by apical dominance. To confirm this hypothesis that multiple shoot development was more efficient from the cot-node tissue, SAMs were purposely removed by cutting through the middle of each epicotyl (decapitated EA explants) (Figure 1a and e) and the isolated EA was cultured for regeneration on SIM. Indeed, removal of the SAM purposely by cutting through the middle of each epicotyl (decapitated EA explants) (Fig. 1a and e) induced 78% of explants to initiate multiple shoot regeneration in IT86D-1010 (Figure 1f, g and Table 1).

**Figure 1.**
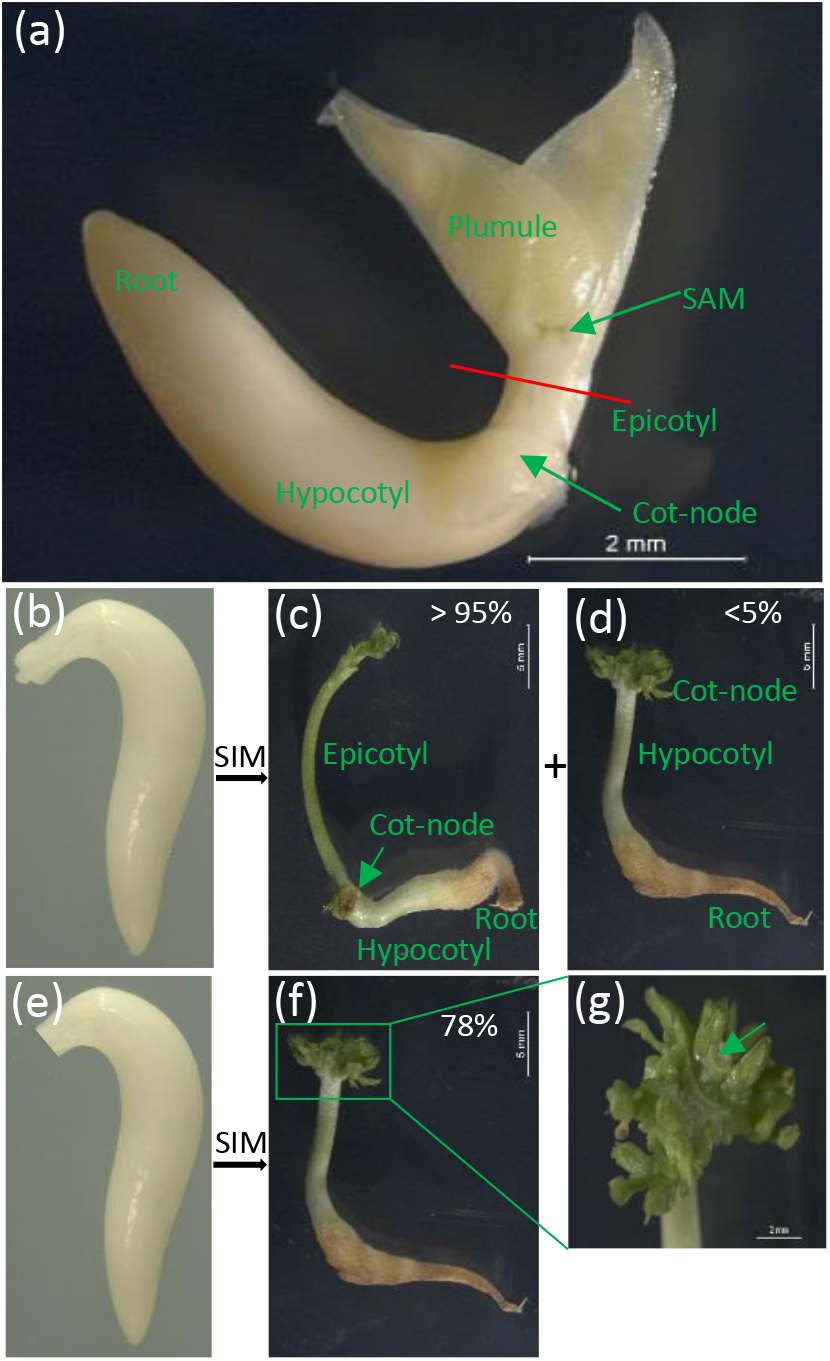
The regeneration origin of cowpea EA explants *via* organogenesis. (a) The structure of cowpea IT86D-1010 EA extracted from imbibed mature seed. The red line through the middle of epicotyl represents the decapitation process of EA explant to remove the SAM. (b-d) The single (c) and multiple (d) shoot development from non-decapitated EA explants (b). (e-f) The multiple shoot regeneration (f) from decapitated EA explants (e). (g) Close look of multiple shoot regeneration and the cutting site of the epicotyl indicated by the arrow. The percentages in pictures (c, d and f) represent the population of single and multiple shoot development determined from 100 non-decapitated and 100 decapitated EA explants after 10 days on SIM.

**Table 1.** Shoot organogenesis of selected accessions of cowpea and common bean

| Germplasm accessions | # of EAs | # of EAs with single shoot (%) | # of EAs with multiple shoots (%) | Explants forming shoots (%) |
| --- | --- | --- | --- | --- |
| *Cowpea* |  |  |  |  |
| IT86D-1010 | 36 | 0 (0) | 28 (78) | 78 |
| PI 527675 | 36 | 0 (0) | 29 (81) | 81 |
| PI 580227 | 36 | 2 (6) | 26 (72) | 78 |
| PI 582835 | 29 | 1 (3) | 22 (76) | 79 |
| PI 583259 | 36 | 3 (8) | 17 (47) | 56 |
| TVu 8670 | 36 | 2 (6) | 22 (61) | 67 |
| TVu 3562 | 37 | 4 (11) | 21 (57) | 68 |
| TVu 9693 | 36 | 0 (0) | 22 (61) | 61 |
| TVu 79 | 36 | 1 (3) | 28 (78) | 81 |
| *Mexican cowpea* |  |  |  |  |
| TPC-001 | 50 | 6 (12) | 12 (24) | 36 |
| MRS-001 | 50 | 5 (10) | 14 (28) | 38 |
| *Common bean* |  |  |  |  |
| CBB-001 | 50 | 4 (8) | 11 (22) | 30 |
| CBP-001 | 50 | 6 (12) | 8 (16) | 30 |

To test if the regeneration principle described above was applicable to other cowpea germplasm accessions and even common bean, we further tested tissue culture and *in vitro* regeneration procedures for eight additional cowpea accessions from the U.S. National Plant Germplasm System (NPGS), two non-conventional cowpea germplasm lines (TPC-001 and MRS-001) and two common bean (*Phaseolus vulgaris* L) varieties, *black* bean (CBB-001) and *pinto* bean (CBP-001), collected from tropical and subtropical regions of Mexico (Figure S3). Consistent with observations in IT86D-1010, shoot regeneration also developed exclusively from cot-node regions for all ten cowpea germplasm lines and two bean germplasm lines tested (Figure S4). In most cases, the number of EA explants showing multiple shoot regeneration exceeded those showing a single regenerated shoot, suggesting that multiple individuals can be recovered from a single EA explant (Table 1 and Figure S4). The overall regeneration efficiency ranged from 55% to 81% for eight additional cowpea accessions from the NPGS, 36 to 38% for Mexican cowpeas and 30% for common beans (Table 1). These results demonstrated that tissue culture and regeneration procedures can be applied to a wide collection of cowpea germplasm and extended to other legumes such as common bean.

### Regeneration optimization under Agrobacterium-mediated transformation

Generally, the EA-based dicot transformation procedure consists of the following main steps: explant preparation, *Agrobacterium* infection, co-cultivation, shoot regeneration with selection and root induction (Figure S2. Also see Experimental Procedures for detail). As described above, although EA explants per se have been described for soybean transformation, the cowpea EA-based regeneration system has a key difference. While the soybean EA transformation system relies on preexisting apical meristematic tissue for regeneration and transformation (Aragão *et al*., 2000; Liu *et al*., 2004; Turlapati *et al*., 2008), the cowpea system shows that removal of the SAM stimulated multiple shoot organogenesis from the cot-node. This key difference raises the question of how well the decapitated-EA explants will be able to survive and regenerate throughout the *Agrobacterium*-mediated transformation procedure.

To evaluate how the decapitation of EA explants affects the survival and regeneration capability, we conducted sonication, *Agrobacterium* infection and co-cultivation treatments, (Figure S2. Also see Experimental Procedures for detail) either with or without *Agrobacterium*, followed by regeneration on SIM. This allowed us to measure the survival and regeneration of the decapitated and non-decapitated EA explants and assess possible damage caused by the early steps of transformation, such as sonication-associated wounding, potential medium component toxicity, sensitivity to *Agrobacterium* and the length of the treatment steps before regeneration. As shown in Figure 2a, the decapitated EA explants were extremely sensitive to the treatments and none of the EA explants survived on SIM without selection after mimicking all the treatment steps without *Agrobacterium* infection. On the contrary, all the non-decapitated EA explants survived and formed elongated epicotyls with a single primary shoot. The further decapitation of those primary shoots by cutting through the middle of the elongated epicotyls after 4-days of regeneration stimulated multiple shoot organogenesis around the cot-nodes (Figure 2b). Collectively, those observations suggest that although the SAM negatively regulates multiple shoot organogenesis from cot-node tissue because of the apical dominance effect, the SAM is essential for EA explant survival through the transformation treatments before regeneration and the SAM should not be removed until fully recovered after 4 days of regeneration.

**Figure 2.**
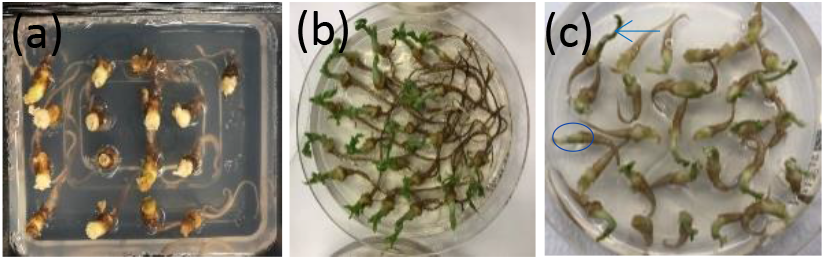
Explant sensitivity to the infection and co-cultivation treatments. (a-b) The survival and regeneration capability of decapitated (a) and non-decapitated (b) EA explants after 10-day shoot induction without selection following infection and co-cultivation steps without *Agrobacterium* inoculation. (c) The susceptibility of non-decapitated EA explants to the *Agrobacterium* inoculation following infection and co-cultivation steps. Image was taken on 4-day SIM. Arrow and circle indicates one of the survived explants with elongated epicotyl and one of the dying explant without elongated epicotyl, respectively.

It has been reported that *Agrobacterium*-mediated infection leads to cell damage and tissue necrosis (Norkunas *et al*., 2018). To determine the survival rate of EA explants after infection and co-cultivation with *Agrobacterium*, non-decapitated EA explants were subjected to the transformation procedure (Figure S2. Also see Experimental Procedures for transformation procedure) using LBA4404 Thy-carrying the pPHP86170/pPHP71539 vector system as described below. As shown in Figure 2c, about 70 ± 10% EA explants (based on the average of 3 replicates and total 75 EA explants) survived and formed elongated epicotyls after 4-day regeneration on SIM with selection (Table S5). Compared to the 100% survival rate without *Agrobacterium* inoculation (Figure 2b), the 30% rate of the explant loss was most likely due to the sensitivity of EA explants to the *Agrobacterium*.

Based on those studies, the decapitation of EA explants after 4 or 5 days of regeneration on SIM was routinely performed for all the subsequent transformation optimization experiments throughout this study.

### Agrobacterium-mediated gene delivery using ternary vector system

A dramatic increase of T-DNA delivery efficiency was reported in cowpea by constitutive expression of additional *vir* genes in a resident pSB1 vector in *Agrobacterium* strain LBA4404 (Solleti *et al*., 2008b). Recently, we demonstrated that a newly designed ternary vector containing the T-DNA binary vector and the optimized pVIR accessory (pPHP71539) plasmid with additional *vir* genes enhanced gene delivery and ultimately the transformation efficiency for both corn and sorghum (Anand *et al*., 2018; Che *et al*., 2018). This encouraged us to assess the gene delivery efficiency of the ternary vector for cowpea transformation.

To evaluate T-DNA delivery using the ternary vector system in cowpea, we transformed binary vector pPHP86170 (Figure S1a) containing the *proDMMV:TagRFP* as visual marker and *proGM-UBQ:CTP-spcN* as the selectable marker into the *Agrobacterium* strain LBA4404 Thy-harboring the pVIR accessory plasmid pPHP71539. Transient gene delivery was assessed by visually evaluating the number of fluorescent foci on the surface of cowpea EA explants after 3 days of shoot induction on SIM containing 25 mg/l spec as selection. As shown in Figure 3b, strong gene delivery based on the number of infected cells was visualized across the entire explant for those surviving EA explants with elongated epicotyls (Figure 3a and b), especially around the cot-node tissue (Figure 3b and c), demonstrating highly efficient gene delivery in cowpea EA explants using the ternary vector system. Although gene delivery was efficient across the entire EA explant, only those fluorescent foci within the cot-node tissue showed subsequent development and substantially enhanced fluorescence intensity during regeneration (Figure 3b and c). This observation supported the hypothesis that only those cells within the cot-node tissue, but not any other tissues of the EA explant, actively undergo rapid cell division and dedifferentiation to acquire organogenetic competence for shoot regeneration.

**Figure 3.**
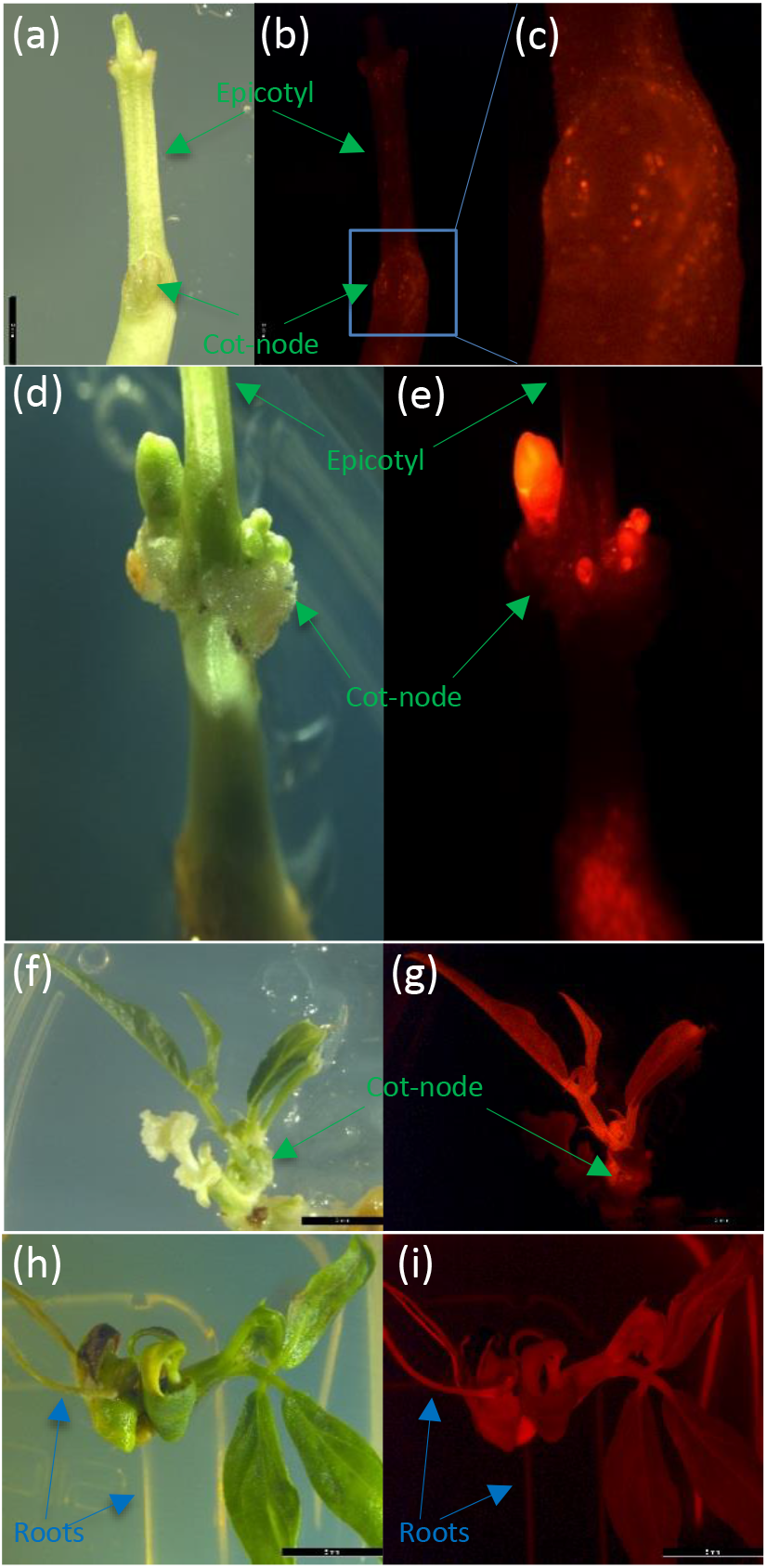
Different stages of transgenic cowpea IT86D-1010 development using *CTP-spcN*/spec selection system. (a-b) T-DNA delivery determined by transient assay after 3-day shoot induction. (c) Close look the number, size and intensity of fluorescent foci around the cot-node region. (d-e) Transgenic shoot budding after 2-week regeneration on SIM with spec selection. (f-g) Fully developed transgenic shoot after 5-week regeneration with spec selection. (h-i) Root development of regenerated transgenic shoot after 3-week root induction on RIM. (a, d, f and h Bright field images. (b, c, e, g and i) Fluorescence images under RFP filter.

### In vitro regeneration and transgenic selection of cowpea

Several selection systems have been reported for cowpea transformation with different explant types (Manman *et al*., 2013), such as *NPTII*/kan (Bett *et al*., 2019; Chaudhury *et al*., 2007), *NPTII*/G418 (Solleti *et al*., 2008b), *PMI*/mannose (Bakshi *et al*., 2012), *HPT*/hygromycin (Kumar *et al*., 1996), *BAR*/glufosinate (Popelka *et al*., 2006) and *ahas*/imazapyr (Citadin *et al*., 2013; Ivo *et al*., 2008). It was reported that incomplete selection and tissue necrosis were associated with those selection systems and resulted in lower transgenic plant recovery in cowpea (Bakshi *et al*., 2011; Chaudhury *et al*., 2007; Solleti *et al*., 2008b). To identify more efficient selection agents for the *de novo* organogenesis described above, we tested and compared the selection efficiency of *CTP-NPTII*/kan, *CTP-NPTII*/G418 and *CTP-spcN*/spec after *Agrobacterium*-mediated transformation using *Agrobacterium* strain LBA4404 Thy-. We found that the utilization of *CTP*-*spcN*/spec system provided more efficient and stringent transgenic selection than either *CTP-NPTII*/kan or *CTP-NPTII*/G418 and without non-transgenic escapes and chimeric tissue formation. As described above, strong gene delivery was observed around the cot-node target tissue on the third day of regeneration as shown in Figure 3b and c. Those fluorescent foci grew quickly and single or multiple shoot buds emerged exclusively around the cot-nodes within 2 weeks following removal of the SAM on the fourth day of regeneration (Figure 3d and e). Transgenic shoots were fully developed from the buds within another 3 weeks in which all the shoots displayed strong fluorescence evenly across the entire regenerated shoot (Figure 3f and g compared to the regenerated shoots from wild-type cowpea IT86D-1010 EA explants that showed no fluorescence at all (Figure S5 a and b). The elongated shoots were excised from the EA explants and transferred to root induction medium (RIM) (Table S6) for root development. Because of the stringent selection during shoot organogenesis, selection was not required for rooting. Approximately 95% (Table 2) of the elongated shoots fully rooted in the RIM (Table S6) within 2-3 weeks and all the regenerated shoots and roots displayed strong fluorescence (Figure 3h and i). The total time from inoculation of the EA explants to transplantation of a fully developed transgenic plantlet in the greenhouse took approximately 2-3 months. The frequency of shoot formation was about 21% for IT86D-1010, of which about 23% of the events were single-copy quality events (Table 2) (see Experimental Procedures for “quality events” definition and event quality determination).

**Table 2.** Transformation efficiency and event quality for IT86D-1010

| # of EAs | Transgenic shoots | Transgenic shoots with root developed | Rooting efficiency (%) | Transformation efficiency (%) | Quality events (%) |
| --- | --- | --- | --- | --- | --- |
| 30 | 7 | 6 | 86 | 20 | 23 |
| 30 | 7 | 7 | 100 | 23 |  |
| 20 | 4 | 4 | 100 | 20 |  |
| Average* |  |  | 95 ± 8 | 21 ± 2 |  |

\*Data were presented as the average ± SD.

Conversely, cowpea IT86D-1010 EA explants showed a high degree of resistance to kan as a selection agent and no selection pressure could be built up by culturing non-transformed EA explants on SIM containing kan at concentrations as high as 600 mg/l. Although, an optimal concentration of G418 at 20 mg/l for selection of transformed shoots was established by culturing non-transformed EA explants on SIM containing different concentrations of G418 (10-40 mg/l), the G418 selection was not as stringent as spec and only chimeric events (Figure S6a and b) were observed using the binary vector pPHP94518 (Figure S1b) containing *proGM-EF1A2:Ds-RED* as a visual marker and *proGM-UBQ:CTP-NPTII* as the selectable marker. Chimera formation was indicated by uneven and partial fluorescence of the regenerated shoot in Figure S6a and b.

### Transgene inheritance in the progeny

To evaluate the inheritance of T-DNA integration events, we selected three independent single-copy quality T0 transgenic events in the IT86D-1010 background (see Experimental Procedures for “quality events” definition and event quality determination) transformed with construct pPHP92782 (Figure S1c) containing the *proGM-EF1A2:Ds-RED* as a visual marker gene and *proGM-UBQ:CTP-spcN* as the selectable marker. Those T0 plants were self-pollinated in the greenhouse and the resultant T1 seeds displayed either red color (similar seed color with *Ds-RED* expression was reported in soybean (Nishizawa *et al*., 2006)) or no visual pigmentation difference from the wild-type as shown in Figure S7a and b, demonstrating the transmission and segregation of the transgene in the progeny. To further determine the segregation ratio, one hundred T1 seeds from each of the three independent events were randomly chosen regardless of seed color and advanced to the T1 generation. The zygosity (homozygous, hemizygous and null) of individual T1 plants was characterized by determining the copy number of the integrated T-DNA based on the assays described in the Experimental Procedures. As shown in Table 3, the segregation pattern of these transgenic events showed typical 1:2:1 and 3:1 Mendelian ratio based on Chi-square test (χ^2^) for all the copy number assays performed (Figure S1c and Table S9). Those results demonstrate that all the transgenic events analyzed possessed stably integrated T-DNA and the T-DNA was faithfully inherited to the next generation.

**Table 3.** Segregation analysis of self-fertilized IT86D-1010 transgenic cowpea plant in the T1 generation

| Event ID | Total plants analyzed | # of Two-copy plants* | # of single-copy plants* | # of null plants* | $P(\chi^2_{1:2:1})$ | $P(\chi^2_{3:1})$ |
| --- | --- | --- | --- | --- | --- | --- |
| 125739938 | 100 | 17 | 54 | 29 | 0.17 (3.52) | 0.36 (0.85) |
| 125739949 | 98 | 28 | 37 | 33 | 0.04 (6.39) | 0.05 (3.93) |
| 125739950 | 99 | 17 | 57 | 25 | 0.17 (3.57) | 0.95 (0.0033) |

\*Copy numbers were determined by both *Ds-RED* and *spcN-SO* qPCR assays.

### Transformability evaluation of different cowpea genotypes

Genotype dependence is a major limitation of regeneration and *Agrobacterium*-mediated transformation for both monocots and dicots (Abdu Sani *et al*., 2015; Che *et al*., 2018; Jia *et al*., 2015; Manman *et al*., 2013). To evaluate the robustness of the protocol and broaden the application of this cowpea transformation technology for different cowpea genotypes, we performed a quick transformability assay to evaluate the formation of fluorescent transgenic shoots after 2 weeks of culture on SIM with selection. This quick transformability assay was conducted using a hypervirulent *Agrobacterium* strain AGL1 carrying RC2717 plasmid with *CTP-aadA* as the selectable marker (Figure S1d). As shown in Table 4, transgenic shoot regeneration frequency (defined as transformability) of IT86D-1010 determined by the quick transformability assay was in a range of 11 to 26% (average 19% ± 7.5%) (Table 4). This was comparable to the 21% ± 2% transformation efficiency described earlier using LBA4404 Thy-carrying the pVIR accessory plasmid for transformation and *CTP-spcN*/spec for selection (Table 2), indicating the reliability of this quick assay for predicating transformation efficiency of different genotypes. The application of this quick transformability assay to eight more cowpea genotypes showed the transformability in a range of 4.5 to 37%. Similar to the observation of transgenic shoot development for IT86D-1010, all the eight cowpea genotypes also formed transgenic shoots exclusively and rapidly at the cot-node region and, in most of the cases, no more than two transgenic shoots per explants were developed (Figure S8). These results demonstrate that the transformation protocol developed for IT86D-1010 described herein is transferable to other genotypes even with an alternative *Agrobacterium*-mediated transformation system.

**Table 4.** Transformability evaluation of nine cowpea accessions

| Germplasm | # of EAs | # of EAs with fluorescent shoots in 2 weeks | Transformability (%) |
| --- | --- | --- | --- |
| *Exp 1* |  |  |  |
| IT86D-1010 | 123 | 14 | 11 |
| PI 527675 | 166 | 52 | 31 |
| PI 580227 | 178 | 10 | 5.6 |
| TVu 9693 | 160 | 35 | 22 |
| *Exp 2* |  |  |  |
| IT86D-1010 | 117 | 22 | 19 |
| PI 583259 | 119 | 31 | 26 |
| TVu 79 | 157 | 7 | 4.5 |
| TVu 8670 | 125 | 11 | 8.8 |
| *Exp 3* |  |  |  |
| IT86D-1010 | 102 | 27 | 26 |
| PI 582835 | 90 | 33 | 37 |
| TVu 3562 | 177 | 32 | 18 |

In general, better shoot organogenesis response tends to produce higher transgenic shoot regeneration frequency, but this is not always the case. As shown in Table 4 and 1, although all five genotypes, IT86D-1010 and PI 527675, PI 580227, PI 582835 and TVu 79, showed very good and comparable shoot organogenesis response in the range of 78 to 81% efficiency, only IT86D-1010, PI 527675 and PI 582835 demonstrated significant high transformability (19% ± 7.5%, 31% and 37%, respectively), but not PI 580227 and TVu 79 (5.6% and 4.5%, respectively). In contrast, all three genotypes, TVu 3562, TVu 9693 and PI 583259, had relatively low shoot organogenesis response (68%, 61% and 56%, respectively) (Table 1), but their transformability was relatively high (18%, 22% and 26%, respectively) (Table 4). Therefore, the transformability of those germplasm lines is determined not only by the shoot organogenesis capability, but also by the combination of the susceptibility to *Agrobacterium*-mediated T-DNA delivery and sensitivity to the *Agrobacterium* infection and sonication related damages.

Taken together, we have developed a rapid, robust, flexible and highly efficient cowpea transformation system using EA as explants. The principles established in this study have the potential to increase the transformation efficiencies of other legume species and, potentially, other dicot crops. With recent progress establishing cowpea genetic and genomic resources (Lonardi *et al*., 2019; Munoz-Amatriain *et al*., 2017; Spriggs *et al*., 2018; Yao *et al*., 2016), we believe that the broad application of this cowpea transformation system will have immediate and far-reaching impact on cowpea research that will improve cowpea productivity.

## Experimental Procedures

### Agrobacterium strain and vectors

Two *Agrobacterium* strains, the auxotrophic strain LBA4404 Thy-and AGL1, were used in this study. *Agrobacterium* auxotrophic strain LBA4404 Thy-was used with the ternary vector transformation system for cowpea IT86D-1010 transformation. The ternary vector system contains the T-DNA binary vector and the optimized pVIR accessory (pPHP71539) plasmid as previously described by Che *et al*. (2018) and Anand *et al* (2018). The T-DNA binary plasmid pPHP86170 (Figure S1a) contains the PUC ORI, the *NPTIII* bacterial selectable marker, the *TagRFP* reporter gene and *spcN* (Anada *et al*., 2017) as plant selectable marker gene. The binary plasmid pPHP94518 (Figure S1b) contains the PVS1 ORI, the spec bacterial selectable marker, the *Ds-RED* reporter gene and *NPTII* as plant selectable marker gene. The binary plasmid pPHP92782 contains the PVS1 ORI, the *NPTIII* bacterial selectable marker, the *Ds-RED* reporter gene, *spcN* (Anada *et al*., 2017) as plant selectable marker gene (Figure S1c). The ternary design was assembled by first mobilizing the accessory plasmid pPHP71539 in the *Agrobacterium* auxotrophic strain LBA4404 Thy-and selected on media supplemented with gentamycin (25mg/l). Subsequently the binary constructs were electroporated into *Agrobacterium* strain LBA4404 Thy-containing the accessory plasmid and recombinant colonies were selected on media supplemented with gentamycin plus either kan for pPHP86170 and pPHP92782 or spec for pPHP94518. All constructs were then subjected to next generation sequencing and sequence confirmation before conducting transformation experiments.

AGL1 carrying RC2717, a modified pCAMBIA vector (Figure S1d), was used for testing transformation on different cowpea germplasm other than IT86D-1010. The T-DNA region of the binary vector RC2717 contained a soybean-codon-optimized *aadA1* gene (Martinell *et al*., 2017) as a selectable marker and a reporter cassette in which a *TdTomato* reporter cassette flanked by two *loxP* sites was placed in a reversed orientation between a soybean-codon-optimized *ZsGreen* gene and the *Arabidopsis Ubiquitin 10* promoter (Figure S1d). The *TdTomato* gene provided a visible fluorescence marker to identify transgenic shoots after transformation. With the use of the freeze-thaw method (Chen *et al*., 1994), the binary vector was introduced into AGL1 and the recombinant colonies were selected on medium containing 100 mg/l kan.

Materials reported in this paper may contain components subject to third party ownership (e.g., *TagRFP* and *Ds-RED*). Transgenic and genome edited materials may be subject to governmental regulations. Availability of materials described in this paper to academic investigators for non-commercial research purposes under an applicable material transfer agreement will be subject to proof of permission from any third-party owners of all or parts of the material and to governmental regulation considerations. Obtaining the applicable permission from such third-party owners will be the responsibility of the requestor. Transgenic materials reported in this paper may only be made available if in full accordance with all applicable governmental regulations.

### Plant materials and growth conditions

Cowpea varieties IT86D-1010, PI 527675, PI 580227, PI 582835, PI583259, TVu 8670, TVu 3562, TVu 9693, and TVu 79 originally obtained from the U.S. NPGS (https://www.ars-grin.gov/npgs/) were used for this study. Mexican cowpea accessions TPC1-001 and MRS-001 were collected from local farming communities of Tabasco and Morelos States, respectively; common bean (*Phaseolus vulgaris* L) varieties *black* bean (CBB-001) and *pinto* bean (CBP-001) were obtained from local producers in Guanajuato State. Those varieties were maintained in the greenhouse to collect mature seeds for EA explant isolation.

### Cowpea transformation procedure

The main steps of cowpea transformation mediated by *Agrobacterium* were illustrated in Figure S2. The detailed cowpea transformation protocol including *Agrobacterium* preparation, cowpea EA isolation, transformation procedure and medium preparation were described in the Supporting information.

### Microscopy and imaging

Images were taken using a dissecting Leica M165 FC stereo-epifluorescence microscope, with RFP and Ds-RED filters for detection of fluorescence, using the PLANAPO 1.0× objective, 0.63× zoom, and Leica Application Suite V4.7 acquisition software. The autofluorescence of the wild-type regenerated cowpea was evaluated using the same system. For testing transformation on the eight additional cowpea germplasm lines from NPGS, transgenic shoots expressing TdTomato were monitored with a Stemi SVII dissection stereoscope equipped with a HBO illuminator (Carl Zeiss, Thornwood, NY) and a Ds-RED filter (excitation: 545/25 nm, emission: 605/70 nm, Chroma Technology, Bellows Falls, VT). Images were taken by using an AxioCam camera (Carl Zeiss, Oberkochen, Germany) and the AxioVision LE64 software, and composed by using Photoshop CC (Adobe, San Jose, CA).

### Evaluation of transgenic plants

The integrated copy number of the T-DNA of the binary vector in the transgenic plants was determined by a series of Quantitative PCR (qPCR) analyses based on the method previously described by Wu *et al*. (2014). In this study, an endogenous control qPCR assay (*LBS*) (Table S9) was developed using a house-keeping gene annotated as *3-isopropylmalate dehydrogenase* (Vigun05g298700) in the leucine biosynthetic pathway from cowpea (Misra *et al*., 2017). Four qPCR assays (*PINII_TERM, spcN_SO, CTP and UBQ14_TERM*) for pPHP86170 (Figure S1a and Table S9) and two qPCR assays (*Ds-RED, spcN_SO*) for pPHP92782 (Figure S1c and Table S9) were developed to determine T-DNA copy number by normalizing with endogenous control *LBS* assay.

Outside the border integration sites, PCR backbone-specific assays were developed to check for any border read-through (Wu *et al*., 2014). The presence and absence of *Agrobacterium* vector backbone integration of the binary vector was detected based on screening for sequences from three regions outside of the T-DNA integration sites for each vector, such as *SPC*, *LEFTBORDER* and *NPTIII* for plasmid pPHP92782 and *HYG*, *VIRG* and *HYGROMYCIN* for plasmid pPHP86170 (Figure S1a, c and Table S9).

Stable T-DNA integration was confirmed by copy number determination using genomic DNA extracted from the putative T0 transgenic events. The T0 transgenic plants carrying a single-copy of the intact T-DNA integrations without vector backbone for all assays described were defined as quality events (Anand *et al*., 2018; Che *et al*., 2018; Zhi *et al*., 2015). The percentage of quality events was divided by the total number of events analyzed to calculate the quality event frequency. Only quality events were advanced to the greenhouse for the next generation. The zygosity of the T1 plants was established by determining the copy number of the T-DNA for all the event quality assays (Figure S1a, c and Table S9). Chi-square analysis (χ^2^) was performed to determine whether the difference between the observed and expected ratio was statistically significant.

## Acknowledgements

We thank the super binary vector construction team from Corteva Agriscience for their support with vector construction and environmental control group for cowpea planting in the greenhouse. We thank Dr. Hyeon-Je Cho from Corteva Agriscience for developing the soybean transformation procedure which provided instrumental knowledge for this study. We also thank Drs. Ray Collier and Michael Petersen from the Wisconsin Crop Innovation Center for constructing the RC2717 vector. The study was funded by a sub-award from the CSIRO to Corteva, UGA and LANGEBIO under the Capturing Heterosis for Smallholder Farmers grant from the Bill and Melinda Gates Foundation (BMGF). Corteva Agriscience provided funding and in-kind donations for this project as well.

## Author contributions

P.C., S.C., M.S., M.A. and T.J. conceptualized the methods. P.C., S.C., M.S., Z.Z., J.V., P.O., M.A. and T.J. designed research, analyzed the data and wrote the paper. A.S. conducted cowpea transformation and collected the data for IT86D-1010. Z.Z. Y.G. and K.M. conducted transformation and genotyping for other cowpea genotypes collected from NPGS. J.O. and D.O. developed event quality assays. M.T. conducted regeneration experiment for Mexican cowpeas and beans.

## Supporting information

Additional Supporting Information may be found online in the supporting information tab for this article:

## Cowpea transformation protocol

**Figure S1** Schematic representation of the molecular components of constructs used in this study.

**Figure S2** Flow diagram of the cowpea EA-based *Agrobacterium*-mediated transformation process.

**Figure S3** Dry mature seeds of selected accessions of cowpea and common bean.

**Figure S4** Shoot organogenesis of selected accessions of cowpea and common bean.

**Figure S5** Autofluorescence evaluation. (a) The bright field image and (b) fluorescence image under RFP filter of regenerated wild-type cowpea IT86D-1010. The arrow indicates the regenerated roots.

**Figure S6** Development of chimeric event using *CTP-NPTII*/G418 selection system. (a) Bright field image. (b) Fluorescence image under RFP filter.

**Figure S7** Transgene segregation in the progeny. (a) Mature wild-type cowpea IT86D-1010 seeds. (b) Segregated T1 seeds in IT86D-1010 background harvested from T0 plant containing the *proGM-EF1A2:Ds-RED* as visual marker.

**Figure S8** Formation of transgenic shoots expressing *TdTomato* on the EA explants of nine cowpea germplasm lines after 14-d culture on SIM, bar = 1 mm.

**Table S1** Master plate medium

**Table S2** Working plate medium

**Table S3** Infection medium (IM)

**Table S4** Bean germination medium (BGM)

**Table S5** Shoot induction medium (SIM)

**Table S6** Root induction medium (RIM)

**Table S7** Shoot elongation medium (SEM)

**Table S8** 0MS

**Table S9** Primers used for event quality assays

