## Supporting information for "Developing a rapid and highly efficient cowpea regeneration and transformation system using embryonic axis explants"

**Cowpea transformation protocol**

Unless otherwise specified, all the chemicals used for media preparation are from Sigma-Aldrich.

*Agrobacterium preparation*

1. **Master plate preparation:** Streak *Agrobacterium* from glycerol stock on the master plate medium (Table S1) containing different antibiotics based on the *Agrobacterium* strains and the constructs that bacterium carries to make master plates. Incubate the master plates at 28°C for 3-4 days. The master plates can be kept in the fridge to make working plates and last for a month.
2. **Working plate preparation:** Streak a working plate on the working plate medium (Table S2) using a loop of bacteria from the master plate prepared above and incubate the working plate at 28°C for overnight or over 20 hrs for LBA4404 Thy- and AGL1, respectively.
3. **Inoculum preparation:** Collect 5-7 full loops of bacteria from the working plate using a sterile loop, suspend bacteria in 30 mL infection medium (IM) (Table S3) with acetosyringone (AS, 1M stock in DMSO protected from light, final 200 µM) and dithiothreitol (DTT, 1 M stock, final 1 mM) freshly added in a sterile 50 mL centrifuge tube, adjust OD to 0.5.

*Cowpea EA explant preparation*

1. **Seed sterilization:** Cowpea seeds were surface sterilized using chlorine gas made by mixing in 3.5 mL of 12N HCl and 100 mL bleach (5.25% sodium hypochlorite) for 16 hrs.

2. **Seed pretreatment:** Soak sterilized cowpea IT89D-1010 seeds in the Bean Germination Medium (BGM) (Table S4) with ~45mL of water added for ~16 hrs. For other cowpea varieties transformed with AGL1 30 mL 0MS medium (Table S8) was used to replace BGM.
3. **EA explants isolation:** Isolate embryo axis (EA) explants by removing the seed coats, cotyledons and plumules and put them into sterile water in a petri dish until infection.

*Cowpea transformation*

1. **Infection:** Remove water from petri dish (as much as possible), add 15 mL inoculum, and 50 µL sterile Poloxamer 188 10% solution. Wrap the plate with parafilm and sonicate (VWR, Motel 50T, 120Volts, 1 A or FS30H, Fisher Scientific) for 3 sec. After sonication, add additional 10 mL inoculum (total 25 mL in petri dish) to the mix and gently shake on a at ~60 rpm for 1.5 hrs at room temperature.
2. **Co-cultivation:** Remove bacterium and transfer EAs to filter paper (VWR Cat No. 28320-020) blotted with 700 µL IM in a 100 x 25mm petri dish. Thirty EAs can be piled up on the paper (2-4 piles per plate). Seal plates with micropore tape and keep plates in 21$^{o}$C, 45% RH, 7.7 lums/ft$^2$ chamber for 2 days.

*Cowpea regeneration*

1. **Regeneration with selection:** Insert the roots of EAs vertically into SIM (Table S5) with cot-node and SAM above the medium. Incubate the EAs on SIM at 26°C under 24 hrs light conditions. Remove the SAM by cutting through the middle of epicotyl after 4-5 days culturing on SIM to promote axillary shoots formation at the cotyledon node region.

2. **Rooting:** After 3-5 weeks regeneration, harvest shoots bigger than 3 cm by cutting at base of shoot, and place into root induction medium (RIM) (Table S6).
3. **Shoot elongation:** After 3-5 weeks regeneration, if shoots did not reach to 3 cm, they were transferred to the shoot elongation medium (SEM) (Table S7) for 2-4 weeks culture before transfer to RIM.

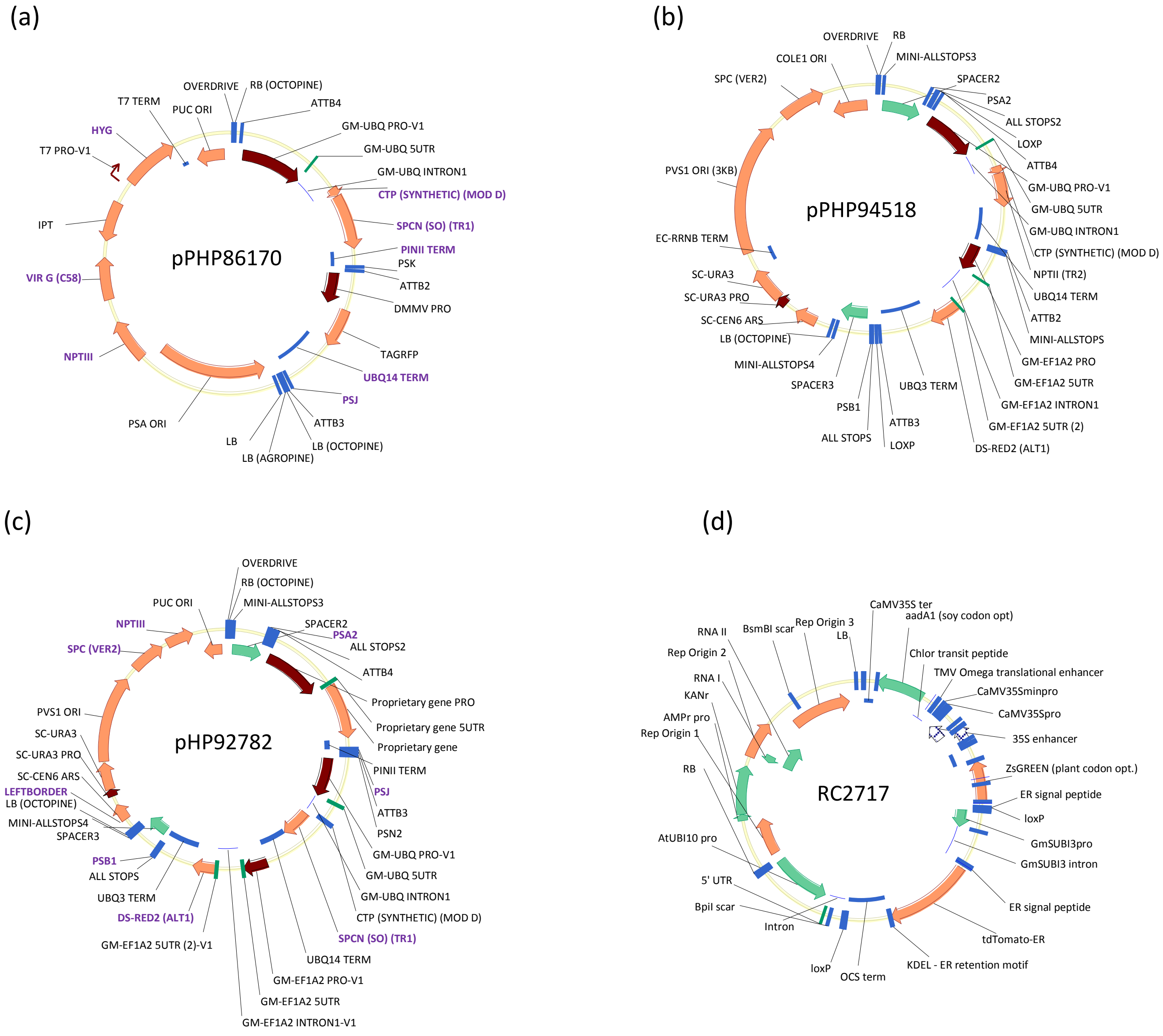


**Figure S1** Schematic representation of the molecular components of constructs used in this study. (a) pPHP86170 (b) pPHP84518 and (c) pPHP92782 were transformed with ternary vector system using pPHP71539 as helper in *Agrobacterium* strain LBA4404 Thy-. The genes and elements highlighted in purple are covered by event quality assays (Table S9). (d) RC2717 is a modified pCAMBIA vector transformed into *Agrobacterium* AGL1 strain.

Seed sterilization and imbibition

↓

EA explants isolation and plumula removal

↓

Sonication in infection medium for 3s and then inoculate with *Agrobacterium* (OD=0.5) for 1.5 hours at room temperature

↓

Co-cultivation on filter paper with infection medium at $21^{o}$C for 2 days

↓

Shoot regeneration on SIM with spec selection for 3-5 weeks following decapitation of EA explants after 4-day regeneration

↓

Root regeneration on RIM for 3 weeks

↓

Rooted plants transferred to greenhouse

**Figure S2** Flow diagram of the cowpea EA-based *Agrobacterium*-mediated transformation process.

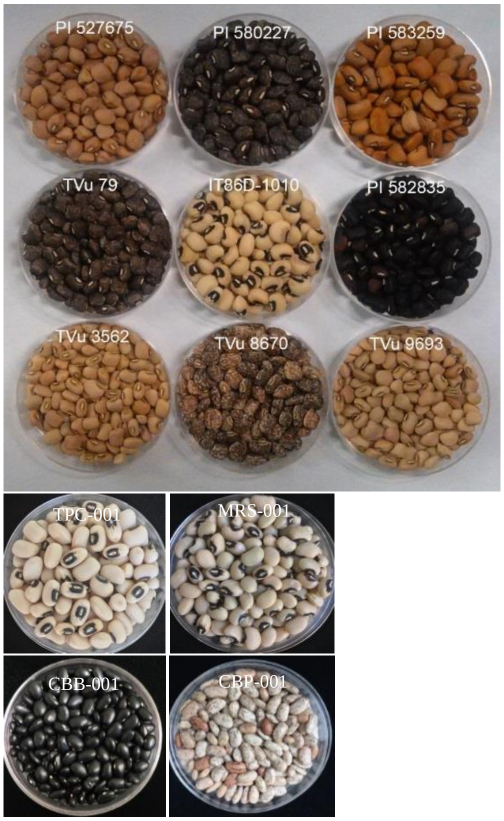


**Figure S3** Dry mature seeds of selected accessions of cowpea and common bean**.**

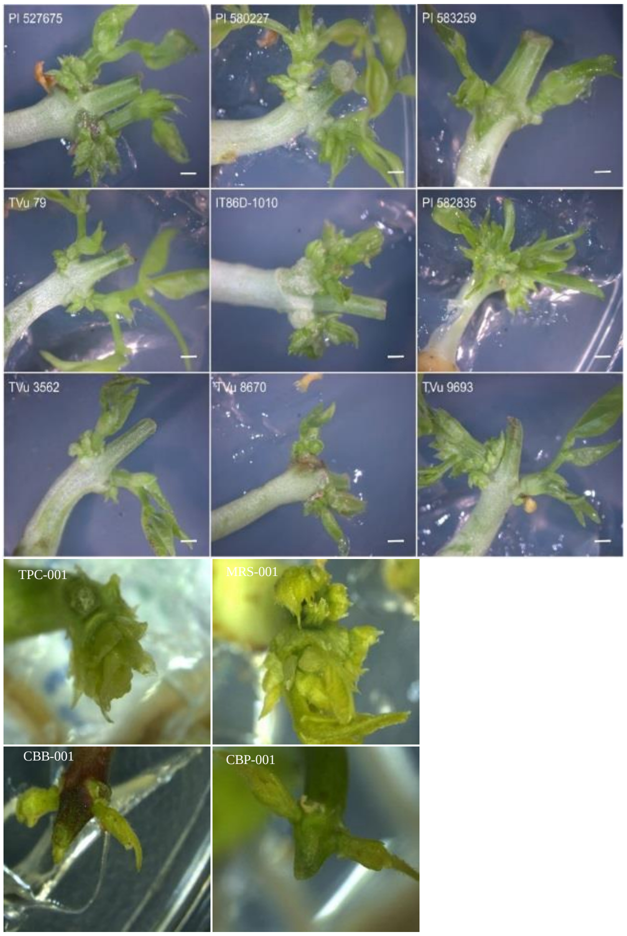


**Figure S4** Shoot organogenesis of selected accessions of cowpea and common bean**.**

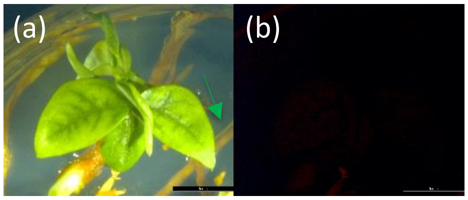


**Figure S5** Autofluorescence evaluation. (a) The bright field image and (b) fluorescence image under RFP filter of regenerated wild-type cowpea IT86D-1010. The arrow indicates the regenerated roots.

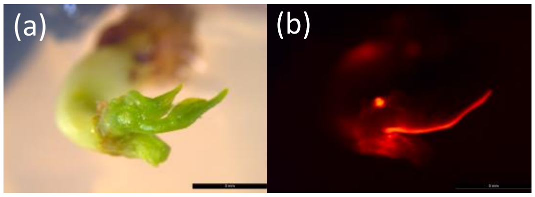


**Figure S6** Development of chimeric event using *CTP-NPTII*/G418 selection system. (a) Bright field image. (b) Fluorescence image under RFP filter.

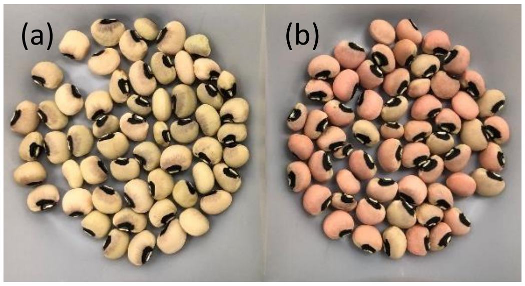


**Figure S7** Transgene segregation in the progeny. (a) Mature wild-type cowpea IT86D-1010 seeds. (b) Segregated T1 seeds (Event ID 125739950) in IT86D-1010 background harvested from T0 plant containing the *proGM-EF1A2:Ds-RED* as visual marker.

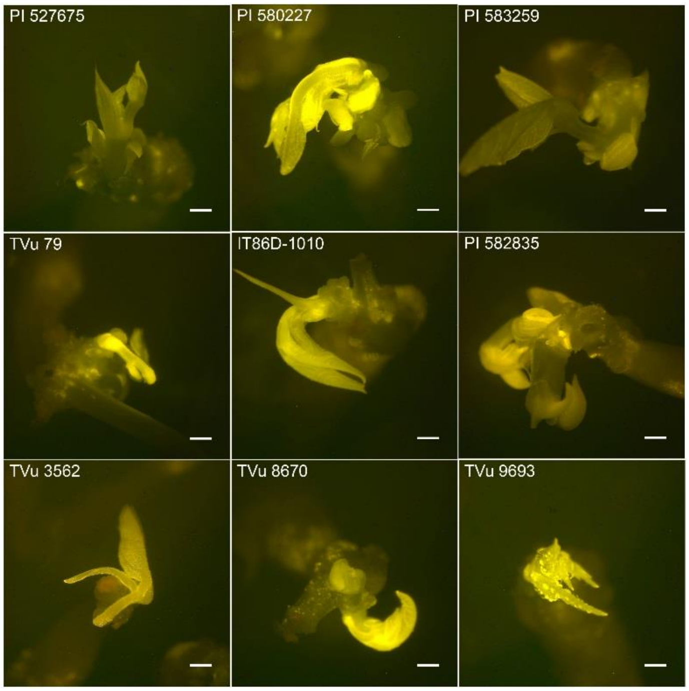


**Figure S8** Formation of transgenic shoots expressing *TdTomato* on the EA explants of nine cowpea germplasm lines after 14-d culture on SIM, bar = 1 mm.

### **Table S1** Master plate medium

| | |
|---|---|
| Glucose | 5 g/l |
| Bacto agar | 15 g/l |
| Ferrous sulfate heptahydrate | 2.5 mg/l |
| Potassium phosphate dibasic | 3 g/l |
| Sodium phosphate, monobasic | 1 g/l |
| Ammonium chloride | 1 g/l |
| Magnesium sulfate heptahydrate | 0.3 g/l |
| Potassium chloride | 0.15 g/l |
| Calcium chloride dihydrate | 11.4 mg/l |
| Thymidine (only for LBA4404 THY- strain) | 50 mg/l |
| Antibiotics* | |

*****100 mg/l kanamycin, 100 mg/l carbenicillin and 25 mg/l rifampicin were used for AGL1 carrying RC2717.
50mg/l gentamycin and 50 mg/l kanamycin were used for LBA4404 Thy- carrying pPHP86170/pPHP71539.
50mg/l gentamycin and 50 mg/l spectinomycin were used for LBA4404 Thy- carrying pPHP94518/pPHP71539 and pPHP92782/ pPHP71539.

### **Table S2** Working plate medium

| | |
|---|---|
| Yeast extract (BD DIFCO) | 5 g/l |
| Peptone | 10 g/l |
| Sodium chloride | 5 g/l |
| Bacto agar | 15 g/l |
| Thymidine (only for LBA4404 THY- strain) | 50 mg/l |
| Antibiotics* | |

*****100 mg/l kanamycin, 100 mg/l carbenicillin and 25 mg/l rifampicin were used for AGL1 carrying RC2717.
50mg/l gentamycin and 50 mg/l kanamycin were used for LBA4404 Thy- carrying pPHP86170/pPHP71539.
50mg/l gentamycin and 50 mg/l spectinomycin were used for LBA4404 Thy- carrying pPHP94518/pPHP71539 and pPHP92782/ pPHP71539.

### **Table S3** Infection medium (IM)

| | |
|---|---|
| MS salt | 1 x |
| MS vitamins | 1 x |
| MES | 20 Mm (3.9 g/l) |
| Sucrose (w/v) | 30 g/l |
| pH | 5.4 |
| BAP | 0.5 mg/l |
| Kinetin | 0.5 mg/l |
| GA3 | 0.25 mg/l |
| Acetosyringone | Add fresh |
| L-cysteine | 400 mg/l |
| BCDA (bathocuproinedisulfonic acid disodium salt) (filter sterilized stock) | 800 µL stock |
| Thymidine (only for LBA4404 THY-) | 50 mg/l |
| Polyvinylpyrrolidone (PVP40) | 1 mg/l |
| Acetosyringone (stock 1M; final 200 µM) | 0.2 mL |
| Dithiothrietol (DTT, stock 1M, filter sterilized aliquot and stored at -80°c; final 1Mm) | 1 mL |

### **Table S4** Bean germination medium (BGM)

| | |
|---|---|
| Sucrose | 25 g/l |
| Thiamine Hydrochloride | 1.34 mg/l |
| Nicotinic acid | 0.5 mg/l |
| Pyridoxine Hydrochloride | 0.82 mg/l |
| EDTA disodium | 3.348 mg/l |

| | |
|---|---|
| Ferrous sulfate heptahydrate | 2.502 mg/l |
| Boric acid | 1.86 mg/l |
| Manganese sulfate, monohydrate | 5.07 mg/l |
| Zinc sulfate, heptahydrate | 2.58 mg/l |
| Potassium iodide | 0.249 mg/l |
| Sodium molybdate dihydrate | 0.216 mg/l |
| Cupric sulfate pentahydrate | 0.00075 mg/l |
| Cobalt chloride hexahydrate | 0.00075 mg/l |
| Calcium chloride dihydrate | 0.176 g/l |
| Potassium nitrate | 0.505 g/l |
| Ammonium nitrate | 0.24 g/l |
| Potassium phosphate monobasic anhydrous | 0.027 g/l |
| Magnesium sulfate heptahydrate | 0.493 g/l |
| TC agar (phytotechnology A296) | 6 g/l |

### **Table S5** Shoot induction medium (SIM)

| | |
|---|---|
| MS salt | 1 x |
| MS vitamins | 1 x |
| MES | 3 Mm (0.59 g/l) |
| Sucrose (w/v) | 30 g/l |
| Agar, DIFCO (w/v) | 8 g/l |
| pH | 5.6 |
| BAP | 0.5 mg/l |
| Kinetin | 0.5 mg/l |
| Antibiotic selection* | |
| Polyvinylpyrrolidone | 1 mg/l |
| Silver nitrate | 2 mg/l |

*25 mg/l spectinomycin and 15 mg/l meropenem was used for EA explants which transformation was carried out through LBA4404 Thy- strain-mediated transformation.

50 mg/l spectinomycin and 15 mg/l meropenem was used for EA explants which transformation was carried out through AGL1 strain-mediated transformation for the first 2 weeks culture and 50 mg/l spectinomycin and 30 mg/l meropenem after the first 2 weeks culture.

### **Table S6** Rooting induction medium (RIM)

| | |
|---|---|
| MS salt | 1 x |
| MS vitamins | 1 x |
| MES | 3 Mm (0.59 g/l) |
| Sucrose (w/v) | 30 g/l |
| Agar, DIFCO (w/v) | 8 g/l |
| pH | 5.6 |
| IBA | 0.1 mg/l |
| Antibiotic selection* | |
| Polyvinylpyrrolidone | 1 mg/l |
| Sliver nitrate | 2 mg/l |

*50 mg/l spectinomycin and 30 mg/l meropenem used for transgenic shoots which transformation were carried out through AGL1 strain-mediated transformation after the first 2 weeks culture.

### **Table S7** Shoot elongation medium (SEM)

| | |
|---|---|
| MS salts | 1 x |
| MS vitamins | 1 x |
| MES | 3 mM (0.59 g/l) |
| Sucrose (w/v) | 30 g/l |

| | |
|---|---|
| Agar (w/v) | 8 g/l |
| pH | 5.6 |
| GA | 0.5 mg/l |
| Kinetin | 0.1 mg/l |
| Asparagine | 50 mg/l |
| Antibiotic selection* | |

*25 mg/l spectinomycin and 15 mg/l meropenem used for transgenic shoots which transformation was carried through LBA4404 Thy- strain-mediated transformation.

50 mg/l spectinomycin and 30 mg/l meropenem used for transgenic shoots which transformation was carried out by AGL1 strain-mediated transformation.

### **Table S8** 0MS

| | |
|---|---|
| MS salts | 1 x |
| MS vitamins | 1 x |
| MES | 3 Mm (0.59 g/l) |
| Sucrose (w/v) | 30 g/l |
| Agar, DIFCO (w/v) | 8 g/l |
| pH | 5.6 |

### **Table S9** Primers used for event quality assay

| Event quality assay | Assay type | Forward primer | Reverse primer |
|---|---|---|---|
| *LBS* | Endogenous control | CACATACCTCCAGTGAGTTCCCTTA | TCGAAGCATCTACTAACTACAGAAGAATTAA |
| *Ds-RED* | Copy number | AAGTCCATCTACATGGCCAAGAA | TGGGAGGTGATGTCCAGCTT |
| *SPCN* | Copy number | CTGCCCGCAATGCTCTTT | ATTACCACTGGACCGTCACAGA |
| *CTP* | Copy number | TGGCTGCAACTACTCTTACATCTG | TGTAAGTTGAAAGGAGCACTTGGT |
| *UBQ14_TERM* | Copy number | CAGAACCCAGAATCCCTTCATATC | TGACGGCTGGGACTTCTTTG |
| HYGROMYCIN | Vector backbone | CAGCGAGAGCCTGACCTATTG | CAGCGAGAGCCTGACCTATTG |
| VIRG | Vector backbone | TGCTCCGAGACGGTCGAT | CAGGCAGGTCTTGCAACGT |
| SPC | Vector backbone | GCGCTGCCATTCTCCAAAT | ATCATTCCGTGGCGTTATCC |
| LEFTBORDER | Vector backbone | GATCTCGCGGAGGGTAGCA | CGAGGGAGATGATATTTGATCACA |
| NPTIII | Vector backbone | CCGATGTGGATTGCGAAAA | GCTCGCGCGGATCTTTAA |
